# Herbal tablet of *Pueraria tuberosa* water extract suppresses the alloxan induced liver damage and hyperglycemia in rats

**DOI:** 10.1101/671594

**Authors:** Harsh Pandey, Shivani Srivastava, Yamini Bhusan Tripathi

## Abstract

**Aim:** To study the protective response of herbal formulation (tablets) of *Pueraria tuberosa* water extract (PTAB) on alloxan induced rat diabetic model.

**Methodology:** Alloxan (120 mg/kg bw) was injected intraperitonially. Rats were divided into three groups: group 1 as normal, group 2 as diabetic control and group 3 were given PTAB upto 14 days. Blood glucose and liver function tests were done using their respective kits. Hematoxylene and eosin staining was done to evaluate the morphological changes in liver tissues. Through immunohistochemistry, we have checked the protein expression of VEGF, MMP9 and ki67.

**Result:** PTAB significantly decreases blood glucose level in a time dependent manner up to 14 days. As compared to diabetic control, PTAB decreases SGOT, SGPT and alkaline phosphates after 14 days of treatment. In diabetic control, the morphology of liver tissues were found damaged due to deformed hepatocytes and dilated lobules. Most of the hepatocytes after PTAB treatment were comparatively found similar to normal rats tissues, along with dilated blood vessels and normalized liver lobules. In addition to these results, PTAB suppresses the expressions of VEGF and MMP 9.

**Conclusion:** PTAB was found to act as an effective hypoglycemic agent. In addition to this, PTAB also found to inhibit inflammation and apoptosis of liver. Thus, can be taken as a potential drug in liver diseases.

## Introduction

Diabetes mellitus is a disease caused due to disorder of carbohydrate metabolism, results in chronic hyperglycaemia, insulin deficit and resistance.(Young-Hyman and Davis, 2010). Metabolic syndrome includes obesity, hypertension, hyperlipidemia and diabetes, primarily caused by oxidative stress and small grade inflammation. Diabetes considered as one of the five foremost causes of death in the world at present scenario. It would be raised from 171 to 366 miliones during 2020-2030 (Wild et al., 2004).Type 1 diabetes was caused by the high dose administration of alloxan /streptozotocin, that initiates the distruction of autoimmune process, resulting in the damages/apoptosis of the beta cells in islets of langerhans. According to this model, alloxan generates reactive oxygen species (ROS) leading to diabetic condition by the mechanism of glucose auto-oxidation and protein glycosylation (Halliwell and Gutteridge, 2003)(Lenzen, 2008). Insulin deficiency leads to various types of metabolic adverse effect, such as increase in blood glucose, alkaline phosphatase and transaminase (Shanmugasundaram etal,1983)(Begun N,1978).The resultant metabolic derangement affects the liver. It leads to elevation in serum bilirubin as well as decrease in serum protein, especially albumin fraction. The liver is associated with the storage of glycogen. The imperative role of the liver in carbohydrate metabolism makes it as an important organ in the pathophysiology of diabetic mellitus..

In this research, we have focused on some fundamental patho physiological mechanism with the help of protein expressions involved in proliferation and vascularisation such as VEGF, MMP9, and Ki67 against alloxan induced liver damages. The stages of the wound healing in liver tissue includes homeostasis, downregulation of inflammation, tissue repairing, which includes cell replication and extracellular matrix (ECM) modulation as well as tissue remodelling (Olczyk et al., 2014). Diabetic ulcer suffers from hyperglycaemia, caused by growth factors disorder such as VEGF and its receptor.The signaling pathways inquisitive with endothelial proliferation, immigration, and recruitment of endothelial progenitor cella (EPC) and bone marrow redemption (Simons, 2005). In the earlier studies, the recombinant VEGF has been used in experimental wound diabetic condition, by *in vivo* and *in vitro*(Brem et al., 2009)(Gan et al., 2005).

Systemic oxidative stress is related to the visceral fat accumulation and metabolic syndrome (Fox et al., 2007). Additionally oxidative stress plays an important role in the progression of nonalcoholic steatohepatitis (NASH)(Sakaguchi et al., 2011) Matrix metalloproteinase (MMP) is a zinc ion-dependent enzyme that helps to control the extracellular matrix, cellular migration and tissue remodelling (Nagase and Woessner, n.d.). MMP 1, MMP-2 and MMP-9 plays an important role in the development of liver fibrosis (Hemmann et al., 2007). The MMP-9 activity and expression are related to oxidative stress (Bittner et al., 2010).

Stress condition induces some other pathological conditions like apoptosis of cells. The protein expression of ki67 binds to the nucleus of proliferating cells, exclusively in all cell cycle except in the Go phase (Srivastava et al., 2018a)(Gerlach et al., 1997). The evaluation of ki67 in hepatocytes proliferative index has been recommended as a useful technique for analyzing liver rejuvenation (Ojanguren et al., 1993).

The use of herbal medicinal plant for treating diabetes is the safer substitute in pharmaceuticals, which can transiently lower the blood glucose, prevents heart disease and high blood pressure and also capable of enhancing the antioxidant system, insulin action and secretion (Surveswaran et al., 2007).

In the present study, we have selected the tubers of *Pueraria tuberosa* Linn (Fabaceae), commonly known as, kudzu(Babu .P et al, 2006) of its tablets. In Sanskrit, it is called Swadukanda, Ikshuganha, Kandapalash and in Hindi, it is called Vidarikanda (Ayurvedic Pharmacopeia, 2006). *Pueraria tuberosa* contains the major isoflavonoids such as Puerarin, tuberostan, genistein, daidzin, tuberosin, pterocarpan, puerarone (Maji et al., 2014)(Singh et al., 2013). Also includes major phyto constituents like steroid, triterpenoids, glycoside, carbohydrates, alkaloids, tannins and amino acids(Tripathi and Kohli, 2013). Phyto-constituents are the major focus for clinical trials and drug discovery.

*Pueraria tuberosa* possesses properties like anti-inflammatory (Tripathi et al., 2013), antioxidant(Pandey et al., 2007), nephroprotective (Tripathi et al., 2012),antihypertension (Verma et al., 2012) and anti hypoglycaemic through inhibition of DPP-IV enzyme (Srivastava et al., 2015)(Srivastava et al., 2018b)(Srivastava et al., 2017)(Srivastava et al., 2018c).

Puerarin is a major active ingredient of *Pueraria tuberosa*. Puerarin is responsible for many pharmacological actions of *Pueraria tuberosa*, studied on the cardiovascular system, cerebrovascular system as well as on antidiabetic effect (Wong et al., 2011) (Wu et al., 2013). On the basis of our previous pre-clinical toxicity study, we have found that water extract of *Pueraria tuberosa* is safe under limit dose (500 mg/ kg bw) and duration (28 days) (Pandey et al., 2018a) and also have prepared its herbal formulation(Pandey et al., 2018b). The tablets of herbal extract of *Pueraria tuberosa*, especially formulation FA has proved good physical properties such as thickness, hardness disintegration and dissolution rate. The starch as an excipients was used at the concentration of 5% or 25 mg for preparation of 500 mg tablets.

Here, we have further evaluated the potency of *Pueraria tuberosa* herbal tablet against liver necrosis induced by alloxan in diabetic rats as well as evaluated the hypoglycaemic response in type 2 diabetic model. Another purpose of this study was to reveal the protective role of tablet through expression study of VEGF, MMP9, and Ki67 in the pathogenesis of diabetes induced liver damage.

## Materials and Methodology

### Reagents and instruments

Alloxan was purchased from Sigma. Polyclonal rabbit VEGF (AR483-5R)-BioGeneX), MMP9, mouse monoclonal KI67 (Biogenex) were the primary required antibodies and the secondary antibody was super sensitive polymer HRP-IHC detection system purchased from biogenesis. Leica RM2125 RT rotator microtome (Leica Biosystem Nussloch GmbH, Nussloch, Germany) and Nikon microscope (Eclipse 50i, loaded with imaging Software –NIS Elements Basic research) were used for imaging.

### Drug preparation

*Pueraria tuberosa* tubers were purchased from Ayurvedic pharmacy, Banaras Hindu University, Ref. no YBT/MC/12/1-2007. Its coarse powder was used to prepare the water extract of *Pueraria tuberosa* (PTWE) by water decoction method..

500 mg tablet (PTAB) was prepared from the obtained extract by wet granulation method. For preparation of suspension we have crushed and dissolved 500 mg of tablet in 2 ml of double distilled water. This suspension was given to rats orally.

### Animals

Charles foster albino male rats were purchased from the central animal house (Ref. 542/GO/ReBi/S/02/CPCSEA) Institute of medical sciences BHU Varanasi. The complete experimental procedure was permitted by our institutional ethical committee (Ref. Dean/2017/CAEC/721).

## Experimental design

Charles foster albino male rats nearly of same age group having weight range of 130-150 g were acclimatized in our laboratory conditions for 7 days with free access to normal standard chow diet and tap water. Before alloxan dose, they were kept in fasting for 8 hours, and then injection was given at the dose of 120mg/kg bw. The rats were then separated into three groups (n=6). viz. group 1 for normal control, 2 for alloxan-induced diabetic control, and group 3 for treatment by PTAB. The animals were sacrificed at their particular days along with normal rats.

### Alloxan Model

Diabetic model was developed by intraperitoneal injection of alloxan monohydrate (120 mg/ kg bw). After five days of alloxan injection, rats with blood glucose levels of >200mg/dl were considered diabetic and used in this study. Treatment of PTAB was started after 5 days of alloxan injection and continued upto next 14 days. Blood samples were collected at 7, 10 and 14 days. After treatment, rats of diabetic control and PTAB group were sacrificed and liver tissues were fixed in 10 % paraformaldehyde for histopathological studies.

## Biochemical evaluation

### Liver function test (LFT) and Blood Glucose

SGOT (Serum glutamic-oxaloacetic transaminase), SGPT (Serum glutamate–pyruvate transaminase), Alkaline Phosphatase (ALP) and plasma glucose were done using their respective Kits, purchased from accurex biomedical Pvt. Ltd.

### Histopathology

Formalin-fixed liver tissue embedded in paraffin was cut into the small sections of 5µm using Leica RM 2125 RT rotator microtome (Leica Biosystems Nussloch GmbH, Germany). The slides were stained with hematoxylin-eosin (H and E). The images were taken with Nikon microscope (Eclipse 50i, loaded with imaging software –NIS Elements Basic research) at 40 X.

### Immunohistochemical assessment

The Poly-L–lysine coated slides were used in immunostaining. Immunostaining for VEGF, MMP-9 and Ki 67 (ready to use) was done according to the kit instructions. Antigen retrieval using citrate buffer, (PH 6.0) was done for 90 min. at 40°C prior to the incubation with primary antibody (Shri SR et al,1991) [38]. After then slides were incubated with secondary antibody “Super sensitive polymer HRP-IHC detection system”. Slides were counterstained with haematoxylin then mounted with DOPEX. Images were taken with light microscope (Nikon) at 40X. Brownish appearance in affected areas is the positive indicator of inflammation.

### Statistical analysis

For each experiment, one-way ANOVA test followed by post hoc analysis with Dunnett’s test was done. all result was expressed as mean±SD. Statistical significant was considered at a p-value less than or equal to 0.05.

## Results

### Liver function test

The SGOT, SGPT, and Alkaline Phosphatase activities were significantly raised in experimental diabetic control animals as compared to normal control. However, PTAB treatment for 14 days significantly decreased this rise in above parameters(Fig 1-B, C, D). **Blood glucose**-The elevated blood glucose level by alloxan were significantly reduced by PTAB, measured at 7, 10 and 14 days. (fig 1 –A).

**Fig 1.**
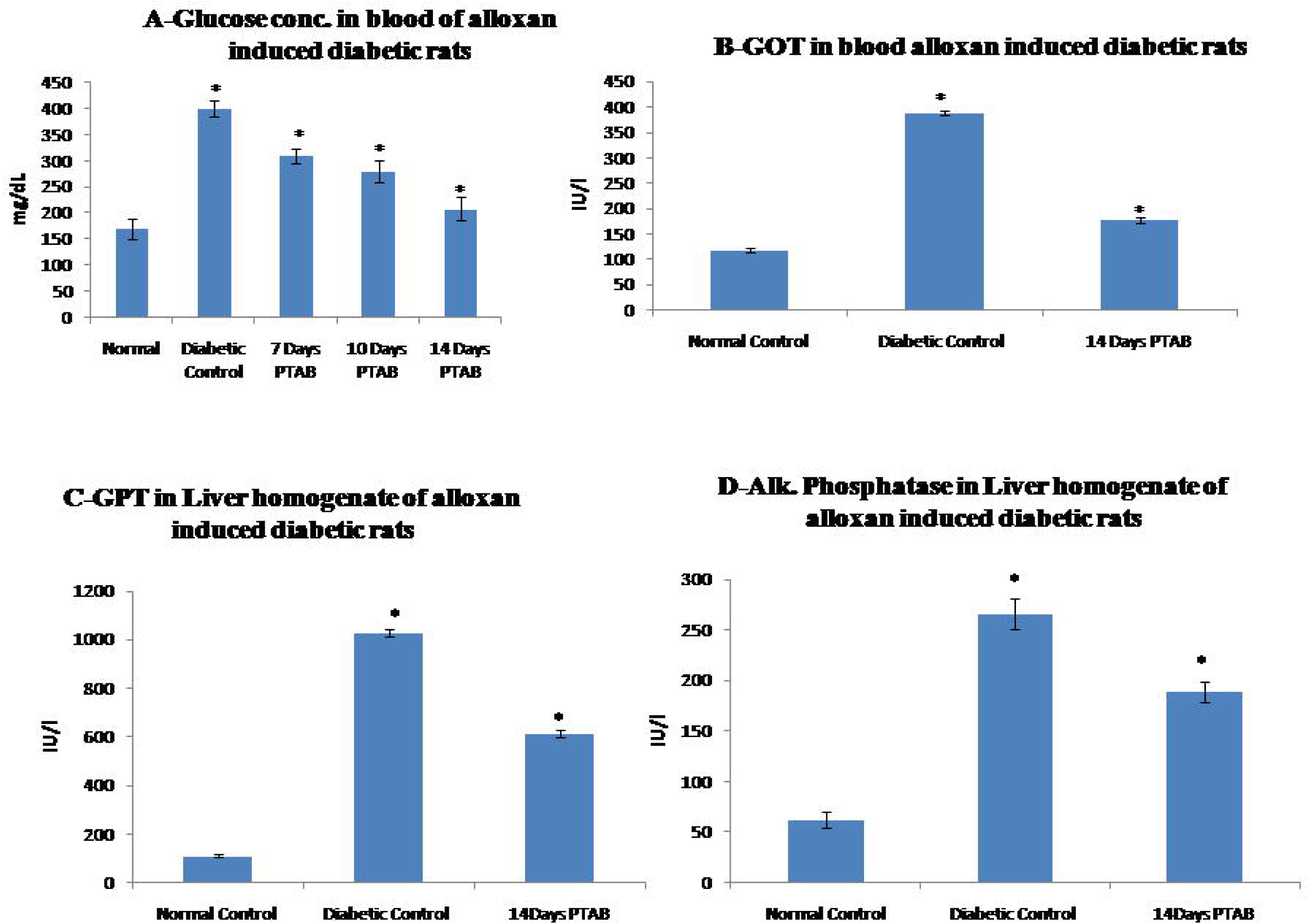
Biochemical analysis: A- Blood glucose of normal, diabetic and 7,10& 14 days PTAB treated group; B-GOT; C-GPT; D-Alkaline phosphatase (ALP) of normal, diabetic and 14 days PTAB treated group.

### Histopathological evaluation

Through microscopic examination, in normal rats, periphery of central vein showed the normal hepatic architecture, also hepatocytes were found in normal morphology. In contrast to this, liver of diabetic rats showed severe injury illustrated by mononuclear cell infiltrate extending through hepatic tissue and hyperplasia. There was severe fibrosis which appears congested with dilation of blood vessels. On the other hand, sections from group third treated with PTAB, showed the curative effect, in which most of the hepatocytes were comparatively similar to normal rats, with dilated blood vessels and normalized liver lobules (fig 2).

**Fig 2.**
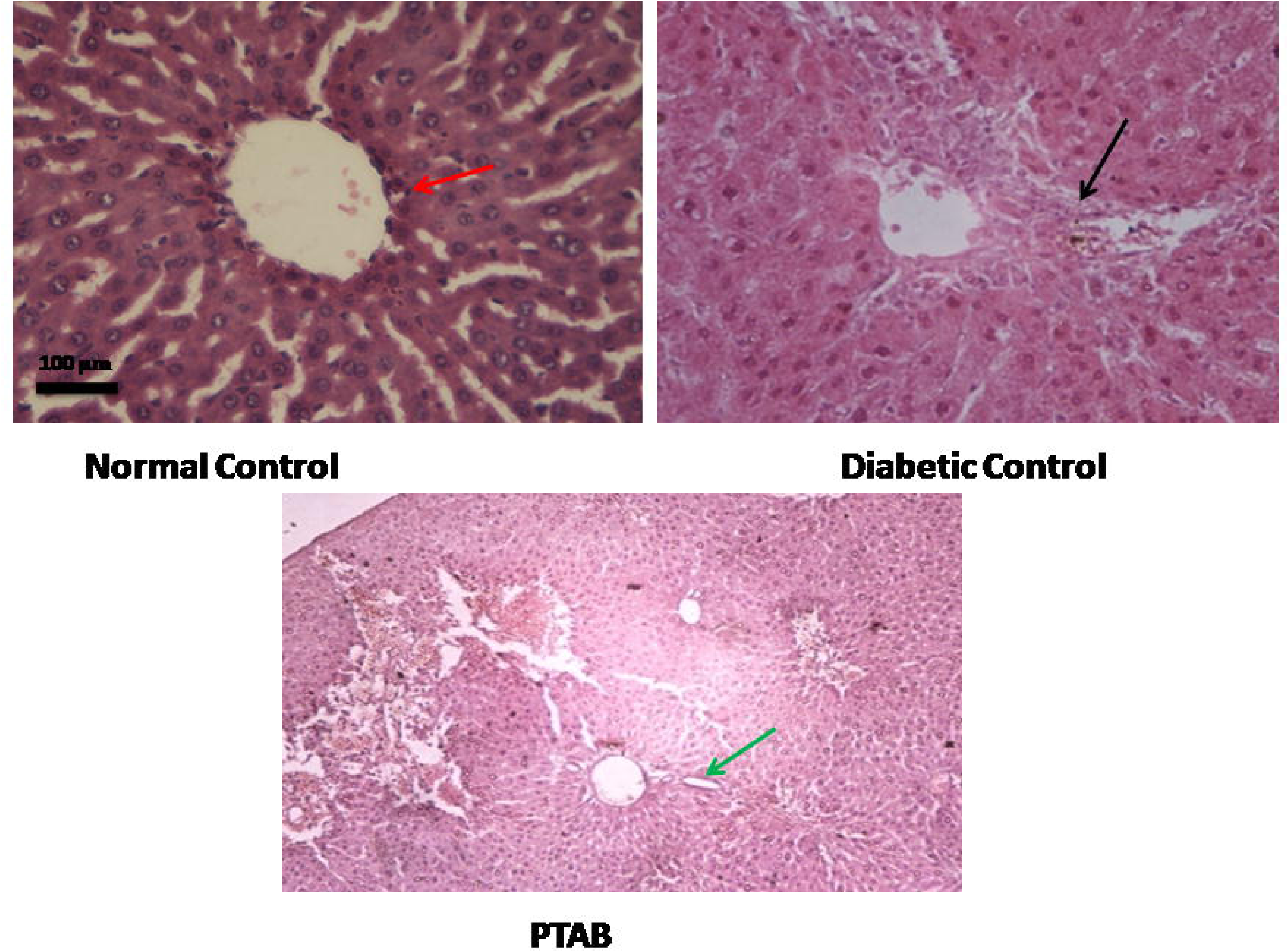
Histopathological analysis of liver: A- Normal; B- Diabetic control; C- PTAB treated group. Black arrow indicates necrosis, red arrow points towards central vein and green arrow is showing hepatocytes regeneration

### Immunohistochemical assessment of VEGF, MMP9 and Ki67 expression

The expression of VEGF and MMP9 significantly decreases in a tissue of PTAB treated group as compared to diabetic control. In contrast to these, Ki67 have shown non-significant changes between each groups.

## Discussion

It is a disease that leads to a serious metabolic disorders affecting human health.. Severe diabetes can causes micro and macro vascular problems (Mohana Lakshmi et al., 2012). Liver diseases like NASH, NAFLD, cirrhosis, acute liver disease etc. are expanding in this era due to changes in lifestyles. The liver is an insulin dependent tissue, which plays an important function in glucose and lipid homeostasis and is severely affected in diabetes. Decrease glycolysis impeded glycogenesis and increase in gluconeogenesis are some of the alterings of glucose metabolism in the diabetic liver (Ivorra et al., 1989).

Many types of allopathic drugs have been developed in this era in order to treat this disease. Due to high cost-effective and many side effects provided by allopathic medicines, many scientists are also doing researches on herbal plants and their products.

This study was undertaken to evaluate the antidiabetic and hepatoprotective activity PTAB in alloxan-induced histopathological alteration. Normal control animal group was found to be stable while the alloxan induced diabetic control group demonstrates high-level cellular abnormalities including, necrosis, hepatocytes infiltration, and hyperplasia. In the PTAB treated group, liver tissues showed hepatocytes regeneration in the area of a central vein. (fig 2).

The elevated activities of GOT, GPT and ALP in blood indicates that diabetes have induced hepatic dysfunction. Here, in our results, PTAB proved as an effective inhibitor of these liver transaminase enzymes (fig 1).

The *Pueraria tuberosa* is a very rich source of flavonoids such as, puerarin, daidzin, genistein, tuberosin and puerarin 4,6-diacetate (Maji et al., 2014)(Singh et al., 2013), those responsible for many pharmacological activities (Asthana et al., 2015).

In this experiment PTAB significantly decreases the blood glucose level in diabetogenic rats (fig 1). The antihyperglycaemic activity caused by Pueraria must be due to the presence of flavonoids (Swanston-Flatt et al., 1990). Puerarin is the major phytoconstituents of tubers of *Pueraria tuberosa*. Puerarin compound has also been reported to be hypoglycaemic against streptozotocin –induced diabetes (Wu et al., 2013).

Alloxan produces different types of reactive oxygen species (ROS). ROS are proficient in oxidizing cellular proteins, nucleic acids, and lipids.

The protective effect of flavonoids found in biologicals are ascribed to their capacity to transfer hydrogen or electron to free radicals(Hanasaki et al., 1994), activates antioxidant enzymes(P.Cos et al 1998), chelates metal catalyst(I.Morel et al 1993), reduces alpha-tocopherol radical (Hirano et al., 2001) and inhibits oxidases (Lima et al., 2014).

VEGF and MMP 9, the pleiotropic proteins generally enhanced in stress-induced diseases and affects many organs like intestine, heart, liver etc. As discussed above, VEGF and MMP9 has a potential role in the development of liver diseases. This tablet preparation could be taken as a treatment against the same.

The circulating VEGF concentration have been reported to be higher in various diseased stage as compare to healthy persons (Ferrari and Scagliotti, 1996). The serum VEGF concentration significantly found to get higher in a diabetic patient as compared to healthy control (Kakizawa et al., 2004). In our case, we have found the protective effect of PTAB, as it reduces the VEGF expression in diabetic liver tissues (Figure 3).

**Fig 3.**
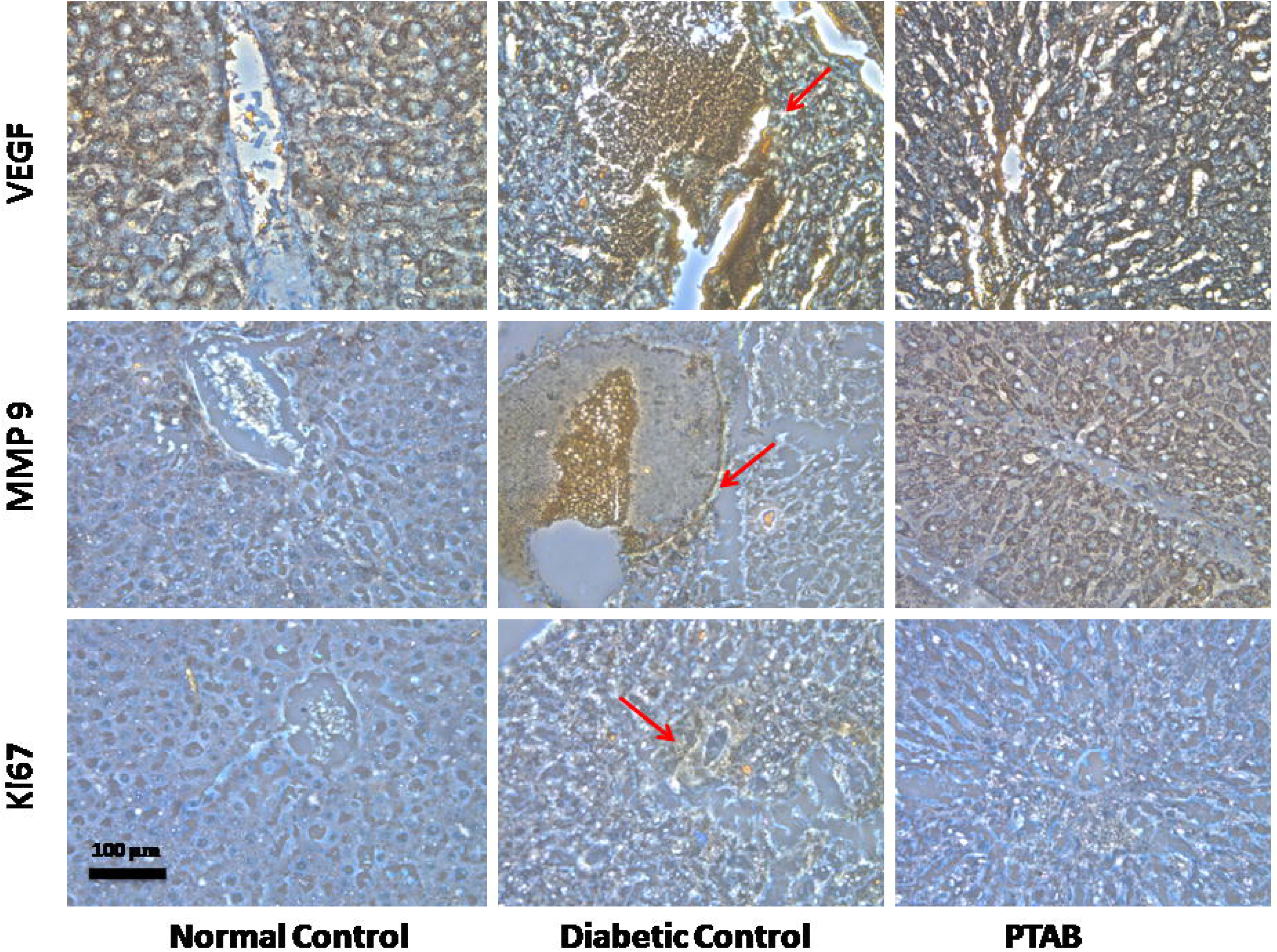
Immunohistochemical staining–. VEGF, MMP9 and Ki67 expression in normal control, diabetic control and PTAB treated liver tissues. Brownish colour indicates relative expression of respective proteins.

MMPs belongs to the proteinase family, that is involved in matrix remodeling and being involved in both physiological and pathophysiological process. The activation of MMPs is sternly controlled by complex formation with tissue inhibitors of MMPs (TIMP). Deregulated expression of both MMPs and TIMPs has been reported in fibrotic liver disease and hepatocellular carcinoma (Iredale, 1997). The effect of MMP9 expression in our diabetic liver tissue was found to be elevated compared to normal tissue, and this elevation were downregulated by PTAB (fig 3)

MMP 9 activity is correlated with endothelial dysfunction and apoptosis in case of diabetes mellitus. The pro-apoptotic role of MMP-9 in hyperglycaemic condition occurs by the activation of caspase -3 (Kowluru and Kanwar, 2009). MMP-9 is formed by kupffer cells (Knittel et al., 1999), induced by IL-beta and TNF alpha. MMP-9 is also involved in tissue inflammation, through ROS, studied in alloxan induced hyperglycemia (Halliwell and Gutteridge, 2003)(Lenzen, 2008).

In contrast to the MMP 9 and VEGF, the expression change of Ki67 was found to be non-significant between the groups. Thus, we can conclude that, here PTAB protects the liver, without involving the proliferative pathway.

This experimental study should be expanded in the future at the cellular and molecular level in order to track the signaling pathways through which PTAB protects the liver by inhibiting the activity VEGF and MMP9.

## Conclusion

PTAB was found to act as an effective hypoglycemic agent against alloxan induced diabetes up to 14 days. In addition to this, PTAB also found to inhibit inflammation and apoptosis as indicated by the reduced expression of VEGF and MMP 9 protein expressions.

## Fundings

UGC -University t fellowship, Goverment of India.

### Acknowledgements

We are thankful to Mr Nikhil Pandey, Ph. D. research scholar of Department of Medicinal Chemistry and Vikas Agrawal, T.A. department of pathology IMS, BHU for his help in sample processing.

## Bibliography

Ayurvedic Pharmacopoeia of India, 2006. Part I, vol. 5, Ministry of Health and Family Welfare, New Delhi, India

Asthana, S., Agarwal, T., Singothu, S., Samal, a, Banerjee, I., Pal, K., Pramanik, K., Ray, S.S., 2015. Molecular Docking and Interactions of Pueraria Tuberosa with Vascular Endothelial Growth Factor Receptors. Indian J. Pharm. Sci. 77, 439–445.

Babu, P.V, Rao, M.A, Kumar, S.M.S, Rao, N.V. 2006. A study on adaptogenic activity of tuber extracts of *Pueraria tuberose*. Indian Drugs. 43(6):486–92

Begum N, Shanmugasudnaram KR. 1978; Tissue phosphates in experimental diabetes, Arogya. J Health Sci.;4:129–39

Bittner, A., Alcaíno, H., Castro, P.F., Pérez, O., Corbalán, R., Troncoso, R., Chiong, M., Mellado, R., Moraga, F., Zanolli, D., Winter, J.L., Zamorano, J.J., Díaz-Araya, G., Lavandero, S., 2010. Matrix metalloproteinase-9 activity is associated to oxidative stress in patients with acute coronary syndrome. Int. J. Cardiol. 143, 98–100. https://doi.org/10.1016/j.ijcard.2008.11.188

Brem, H., Kodra, A., Golinko, M.S., Entero, H., Stojadinovic, O., Wang, V.M., Sheahan, C.M., Weinberg, A.D., Woo, S.L.C., Ehrlich, H.P., Tomic-Canic, M., 2009. Mechanism of sustained release of vascular endothelial growth factor in accelerating experimental diabetic healing. J. Invest. Dermatol. 129, 2275–2287. https://doi.org/10.1038/jid.2009.26

Ferrari, G., Scagliotti, G. V., 1996. Serum and urinary vascular endothelial growth factor levels in non-small cell lung cancer patients. Eur. J. Cancer. https://doi.org/10.1016/S0959-8049(96)00272-9

Fox, C.S., Pou, K.M., Massaro, J.M., Lipinska, I., O’Donnell, C.J., Maurovich-Horvat, P., Kathiresan, S., Larson, M.G., Murabito, J.M., Keaney, J.F., Meigs, J.B., Vasan, R.S., Hoffmann, U., Benjamin, E.J., 2007. Visceral and Subcutaneous Adipose Tissue Volumes Are Cross-Sectionally Related to Markers of Inflammation and Oxidative Stress. Circulation 116, 1234–1241. https://doi.org/10.1161/circulationaha.107.710509

Gan, L.Y., Yu, Z.Y., Cai, M.S., Zhao, H.P., Li, X., 2005. [Effects of low molecular weight heparin on vascular endothelial growth factor expression of early diabetic nephropathy]. Beijing Da Xue Xue Bao 37, 382–385.

Gerlach, C., Golding, M., Larue, L., Alison, M.R., Gerdes, J., 1997. Ki-67 immunoexpression is a robust marker of proliferative cells in the rat. Lab. Invest. https://doi.org/10.1016/j.jaad.2014.11.028

Halliwell, B., Gutteridge, J.M.C., 2003. Free radicals in biology and medicine, second edition. Free Radic. Biol. Med. 10, 449–450. https://doi.org/10.1016/0891-5849(91)90055-8

Hanasaki, Y., Ogawa, S., Fukui, S., 1994. The correlation between active oxygens scavenging and antioxidative effects of flavonoids. Free Radic. Biol. Med. 16, 845–850. https://doi.org/10.1016/0891-5849(94)90202-X

Hemmann, S., Roderfeld, M., Roeb, E., 2007. Expression of MMPs and TIMPs in liver fibrosis – a systematic review with special emphasis on anti-fibrotic strategies 46, 955–975. https://doi.org/10.1016/j.jhep.2007.02.003

Hirano, R., Sasamoto, W., Matsumoto, A., Itakura, H., Igarashi, O., Kondo, K., 2001. Antioxidant ability of various flavonoids against DPPH radicals and LDL oxidation. J. Nutr. … 47, 357–362.

Iredale, J.P., 1997. Tissue inhibitors of metalloproteinases in liver fibrosis. Int. J. Biochem. Cell Biol. https://doi.org/10.1016/S1357-2725(96)00118-5

Ivorra, M.D., Payá, M., Villar, A., 1989. A review of natural products and plants as potential antidiabetic drugs. J. Ethnopharmacol. https://doi.org/10.1016/0378-8741(89)90001-9

Kakizawa, H., Itoh, M., Itoh, Y., Imamura, S., Ishiwata, Y., Matsumoto, T., Yamamoto, K., Kato, T., Ono, Y., Nagata, M., Hayakawa, N., Suzuki, A., Goto, Y., Oda, N., 2004. The Relationship Between Glycemic Control and Plasma Vascular Endothelial Growth Factor and Endothelin-1 Concentration in Diabetic Patients 53, 550–555. https://doi.org/10.1016/j.metabol.2003.12.002

Knittel, T., Mehde, M., Kobold, D., Saile, B., Dinter, C., Ramadori, G., 1999. Expression patterns of matrix metalloproteinases and their inhibitors in parenchymal and non-parenchymal cells of rat liver: Regulation by TNF-α and TGF-β1. J. Hepatol. 30, 48–60. https://doi.org/10.1016/S0168-8278(99)80007-5

Kowluru, R.A., Kanwar, M., 2009. Oxidative stress and the development of diabetic retinopathy: Contributory role of matrix metalloproteinase-2. Free Radic. Biol. Med. 46, 1677–1685. https://doi.org/10.1016/j.freeradbiomed.2009.03.024

Lenzen, S., 2008. The mechanisms of alloxan- and streptozotocin-induced diabetes. Diabetologia. https://doi.org/10.1007/s00125-007-0886-7

Lima, C.C., Lemos, R.P.L., Conserva, L.M., 2014. Dilleniaceae family □: an overview of its ethnomedicinal uses, biological and phytochemical profile. J. Pharmacogn. Phytochem. 3, 181–204.

Maji, A.K., Pandit, S., Banerji, P., Banerjee, D., 2014. Pueraria tuberosa: A review on its phytochemical and therapeutic potential. Nat. Prod. Res. https://doi.org/10.1080/14786419.2014.928291

Mohana Lakshmi, S., Sandhya Rani, K.S., Kiran Reddy, U.T., 2012. A review on diabetes milletus and the herbal plants used for its treatment. Asian J. Pharm. Clin. Res.

Nagase, H., Woessner, J.F., n.d. Matrix Metalloproteinases *.

Ojanguren, I., Ariza, A., Llatjós, M., Castellà, E., Mate, J.L., Navas-Palacios, J.J., 1993. Proliferating cell nuclear antigen expression in normal, regenerative, and neoplastic liver: A fine-needle aspiration cytology and biopsy study. Hum. Pathol. 24, 905–908. https://doi.org/10.1016/0046-8177(93)90141-3

Olczyk, P., Mencner, Ł., Komosinska-Vassev, K., 2014. The Role of the Extracellular Matrix Components in Cutaneous Wound Healing. Biomed Res. Int. 2014, 1–8. https://doi.org/10.1155/2014/747584

Pandey, H., Srivastava, S., Kumar, R., Tripathi, Y.B., 2018a. Preclinical acute and repeated dose toxicity of Pueraria tuberosa (PTWE) on charles foster rats. Int. J. Pharm. Sci. Res. 9, 4572–4581. https://doi.org/10.13040/IJPSR.0975-8232.9(11).1000-10

Pandey, H., Srivastava, S., Mishra, B., Saxena, R., Tripathi, Y.B., 2018b. Development and evaluation of Herbal Tablet loaded with Pueraria tuberosa water extract with use of different Excipients. Asian J. Pharm. 12, 786–793.

Pandey, N., Chaurasia, J.K., Tiwari, O.P., Tripathi, Y.B., 2007. Antioxidant properties of different fractions of tubers from Pueraria tuberosa Linn. Food Chem. 105, 219–222. https://doi.org/10.1016/j.foodchem.2007.03.072

Sakaguchi, S., Takahashi, S., Sasaki, T., Kumagai, T., Nagata, K., 2011. Progression of alcoholic and non-alcoholic steatohepatitis: common metabolic aspects of innate immune system and oxidative stress. Drug Metab. Pharmacokinet. 26, 30–46. https://doi.org/10.2133/dmpk.DMPK-10-RV-087

Shanmugasundaram, K.R., Panneerselvam, C., Samudram, P., Shanmugasundaram, E.R.B., 1983. Enzyme changes and glucose utilisation in diabetic rabbits: the effect of Gymnema sylvestre, R.Br. J. Ethnopharmacol. 7, 205–234. https://doi.org/10.1016/0378-8741(83)90021-1

Simons, M., 2005. Angiogenesis, arteriogenesis, and diabetes: Paradigm reassessed? J. Am. Coll. Cardiol. https://doi.org/10.1016/j.jacc.2005.06.008

Singh, R.R.B., Pandey, M.M., Rawat, A.K.S., Katara, A., Rastogi, S., Arora, S., 2013. Physical Stability and HPLC Analysis of Indian Kudzu (Pueraria tuberosa Linn.) Fortified Milk. Evidence-Based Complement. Altern. Med. 2013, 1–6. https://doi.org/10.1155/2013/368248

Srivastava, S., Koley, T.K., Singh, S.K., Tripathi, Y.B., 2015. The tuber extract of pueraria tuberosa Linn. competitively inhibits DPP-IV activity in normoglycemic rats. Int. J. Pharm. Pharm. Sci. 7, 227–231.

Srivastava, S., Pandey, H., Tripathi, Y.B., 2018a. Expression kinetics reveal the self-adaptive role of β cells during the progression of diabetes. Biomed. Pharmacother. 106, 472–482. https://doi.org/10.1016/j.biopha.2018.06.168

Srivastava, S., Shree, P., Pandey, H., Tripathi, Y.B., 2018b. Incretin hormones receptor signaling plays the key role in antidiabetic potential of PTY-2 against STZ-induced pancreatitis. Biomed. Pharmacother. 97, 330–338. https://doi.org/10.1016/j.biopha.2017.10.071

Srivastava, S., Shree, P., Tripathi, Y.B., 2017. Active phytochemicals of Pueraria tuberosa for DPP-IV inhibition: In silico and experimental approach. J. Diabetes Metab. Disord. 16. https://doi.org/10.1186/s40200-017-0328-0

Srivastava, S., Yadav, D., Bhusan Tripathi, Y., 2018c. DPP-IV Inhibitory Potential of Methanolic Extract of Pueraria Tuberosa in Liver of Alloxan Induced Diabetic Model. Biosci. Biotechnol. Res. Asia 15, 01–04. https://doi.org/10.13005/bbra/2602

Surveswaran, S., Cai, Y.Z., Corke, H., Sun, M., 2007. Systematic evaluation of natural phenolic antioxidants from 133 Indian medicinal plants. Food Chem. 102, 938–953. https://doi.org/10.1016/j.foodchem.2006.06.033

Swanston-Flatt, S.K., Day, C., Bailey, C.J., Flatt, P.R., 1990. Traditional plant treatments for diabetes. Studies in normal and streptozotocin diabetic mice. Diabetologia 33, 462–464. https://doi.org/10.1007/BF00405106

Tripathi, A.K., Kohli, S., 2013. Anti-Diabetic Activity and Phytochemical Screening of Crude Extracts of PuerariaTuberosa DC. (FABACEAE) Grown in India on STZ -Induced Diabetic Rats. Asian J. Med. Pharm. Res 3, 66–73.

Tripathi, Y., Pandey, N., Yadav, D., Pandey, V., 2013. Anti-inflammatory effect of Pueraria tuberosa extracts through improvement in activity of red blood cell anti-oxidant enzymes. AYU (An Int. Q. J. Res. Ayurveda) 34, 297. https://doi.org/10.4103/0974-8520.123131

Tripathi, Y.B., Nagwani, S., Mishra, P., Jha, A., Rai, S.P., 2012. Protective effect of Pueraria tuberosa DC. Embedded biscuit on cisplatin-induced nephrotoxicity in mice. J. Nat. Med. 66, 109–118. https://doi.org/10.1007/s11418-011-0559-1

Verma, S.K., Jain, V., Singh, D.P., 2012. Effect of Pueraria tuberosa DC. (Indian Kudzu) on blood pressure, fibrinolysis and oxidative stress in patients with stage 1 hypertension. Pakistan J. Biol. Sci. 15, 742–747. https://doi.org/10.3923/pjbs.2012.742.747

Wild, S., Roglic, G., Green, A., Sicree, R., King, H., 2004. Global Prevalence of Diabetes: Estimates for the year 2000 and projections for 2030. Diabetes Care 27, 1047–1053. https://doi.org/10.2337/diacare.27.5.1047

Wong, K.H., Li, G.Q., Li, K.M., Razmovski-Naumovski, V., Chan, K., 2011. Kudzu root: Traditional uses and potential medicinal benefits in diabetes and cardiovascular diseases. J. Ethnopharmacol. https://doi.org/10.1016/j.jep.2011.02.001

Wu, K., Liang, T., Duan, X., Xu, L., Zhang, K., Li, R., 2013. Anti-diabetic effects of puerarin, isolated from Pueraria lobata (Willd.), on streptozotocin-diabetogenic mice through promoting insulin expression and ameliorating metabolic function. Food Chem. Toxicol. 60, 341–347. https://doi.org/10.1016/j.fct.2013.07.077

Young-Hyman, D.L., Davis, C.L., 2010. Disordered eating behavior in individuals with diabetes: Importance of context, evaluation, and classification. Diabetes Care. https://doi.org/10.2337/dc08-1077

